# Adaptive deletion of functional duplicate genes in *Drosophila*

**DOI:** 10.64898/2025.12.19.695606

**Authors:** Andre Luiz Campelo dos Santos, Raquel Assis

## Abstract

Gene deletion is traditionally viewed as a nonadaptive mechanism that eliminates functional redundancy, yet emerging evidence indicates that it disproportionately affects tissue-specific duplicates with unique functions. Here, we test whether gene deletion preferentially removes weakly constrained, degenerating duplicates or instead eliminates functionally active duplicates as an adaptive genome streamlining process. To identify the evolutionary and functional factors that determine which duplicates are lost, we systematically analyzed 100 gene deletion events in *Drosophila* by integrating sequence, expression, interaction, and structural data. We uncovered a strong bias toward the loss of younger child copies among functionally unique duplicates, whereas no such bias was observed for redundant duplicates. Contrary to expectations under relaxed constraint, deleted unique genes evolve more slowly, show higher expression, engage in more protein–protein interactions, and do not exhibit elevated structural divergence or intrinsic disorder relative to redundant duplicates. When compared with single-copy genes, deleted unique genes display similar evolutionary rates, slightly lower expression, greater network connectivity, comparable structural divergence, and lower intrinsic disorder. These patterns suggest that deletion frequently targets functionally active rather than degenerate genes. Collectively, our results support the hypothesis that gene deletion in *Drosophila* can represent an adaptive process that removes transiently functional duplicates, promoting genome streamlining and regulatory stability.

**SIGNIFICANCE:** Gene deletion is commonly assumed to be a nonadaptive consequence of redundancy or functional decay after gene duplication. This view has shaped how gene loss is interpreted across evolutionary studies. Using 100 gene deletion events in *Drosophila*, we show that this assumption can be misleading. Deletion strongly favors loss of the younger child copy among functionally unique duplicate genes, whereas redundant duplicates show no such bias. Deleted unique genes also lack signatures of degeneration: they evolve slowly, show relatively high expression, participate in extensive protein–protein interaction networks, exhibit typical levels of structural divergence, and tend to be structurally ordered. Together, these results support a model in which gene deletion can selectively remove transiently functional duplicates, contributing to genome streamlining and regulatory stability. More broadly, our study reframes gene deletion as a potentially adaptive force shaping gene content evolution.

## INTRODUCTION

Gene deletion has long been viewed as a largely nonadaptive process that streamlines the genome by removing redundant duplicate genes (Albalat & Cañestro 2016; Assis 2019). However, mounting evidence shows that deletion often targets duplicates with distinct expression profiles and biological functions (Albalat & Cañestro 2016; Assis 2019; Campelo dos Santos et al. 2024; Hottes et al. 2013; Kvitek & Sherlock 2013), suggesting a potential role in adaptation. The “less-is-more” hypothesis posits that deletion can be beneficial when the function of a gene becomes obsolete, deleterious, or incompatible with newly advantageous traits (Olson 1999). In such cases, deletion may represent an adaptive mechanism for shedding unnecessary or harmful genes— conserving energy, preventing detrimental interactions, or facilitating specialization and niche adaptation (Olson 1999).

A recent study in *Drosophila* revealed that deleted genes are often tissue-specific, with primary expression in testis and accessory gland male reproductive tissues (Campelo dos Santos et al. 2024). Such genes are also expected to evolve rapidly at the sequence level and engage in relatively few protein-protein interactions, reducing the risks of deleterious pleiotropic effects upon their loss (Birchler & Yang 2022; Kuzmin et al. 2022; DeLuna et al. 2010). Intriguingly, these same features are characteristic of recently duplicated genes (Ma et al. 2024; Kaessmann 2010; Kondo et al. 2017; Luis Villanueva-Cañas et al. 2017; Assis & Bachtrog 2013), raising the possibility that deletion preferentially removes the younger “child” copy rather than the ancestral “parent” of a duplicate pair (Han and Hahn 2009; Assis and Bachtrog 2013; Assis 2019). Together, these observations motivate two alternative models: one in which weakly constrained, degenerating duplicates are preferentially lost, and another in which functionally active duplicates are removed as an adaptive mechanism for genome and regulatory streamlining. Discriminating between these models requires systematic comparison of the evolutionary and functional properties of deleted genes.

Here, we address these questions by examining 100 recent gene deletion events in *Drosophila* (Assis 2019; Campelo dos Santos et al. 2024). Each event involves a gene duplication in the common ancestor of *D. melanogaster* and *D. pseudoobscura*, followed by deletion of one copy in a descendant lineage. In these cases, one species represents the derived state after deletion and carries a single “derived” copy (D), whereas the other species represents the ancestral state and harbors two copies: the orthologous “survived” copy (S) and the nonorthologous “lost” copy (L; Campelo dos Santos et al. 2024). In a prior study, we classified these deletion events as targeting either functionally redundant or unique genes (Campelo dos Santos et al. 2024), providing a valuable framework for dissecting the evolutionary dynamics of gene loss. In the present study, we build on that work by asking two central questions: (1) Does deletion preferentially target parent or child duplicate gene copies? and (2) What are the sequence, expression, and protein-level features of deleted genes? By answering these questions in the context of redundant versus unique duplicates, we aim to distinguish between nonadaptive gene loss and adaptive purging of functionally active duplicates, thereby illuminating the evolutionary processes shaping gene deletion in *Drosophila*.

## RESULTS

To determine whether deletion preferentially targets parent or child copies of duplicate gene pairs, we classified parent–child relationships for the 100 gene deletion events in *Drosophila* (Assis 2019). Assignments were based on literature review, sequence conservation across *Drosophila* species, and evidence of retrotransposition (see *Methods*). Using these combined criteria, we identified parent and child copies for 90 of the 100 deletion events. Incorporating functional classifications from Campelo dos Santos et al. (2024), parent–child relationships could be resolved for 37 of 45 redundant deletions and 53 of 55 unique deletions. As expected given their greater expression divergence (Campelo dos Santos et al. 2024), unique genes were significantly more likely than redundant genes to have resolvable parent–child relationships (*p* = 1·97 × 10^−4^, binomial test; see *Methods*).

Among the 37 redundant deletions with resolved relationships, 14 involved deletion of the parent and 23 involved deletion of the child, a difference that did not statistically deviate from the null expectation of equal loss (*p* = 0·19, binomial test; see *Methods*). In contrast, among the 53 unique deletions with resolved relationships, only nine involved deletion of the parent, revealing a strong bias toward child loss (*p* = 1·22 × 10^−6^, binomial test; see *Methods*). Thus, deletion shows no detectable parent–child bias among redundant duplicates, but strongly favors the loss of the child copy among unique duplicates. These results suggest a mechanism for purging recently evolved functions that are no longer beneficial, as child genes in *Drosophila* often acquire novel roles after duplication (Assis & Bachtrog 2013; DeGiorgio & Assis 2021; Kondo et al. 2017; Assis 2014; Jiang & Assis 2017).

To better understand the evolutionary dynamics of gene deletion, we next examined sequence divergence using *K*_a_/*K*_s_ ratios, which compare the rates of nonsynonymous (*K*_a_) and synonymous (*K*_s_) substitutions (Li 1993; see *Methods*). We used single-copy genes as a baseline, comparing distributions of their *K*_a_/*K*_s_ ratios to those of deleted redundant and deleted unique genes. Intriguingly, deleted redundant genes showed significantly elevated sequence evolutionary rates relative to single-copy genes, whereas deleted unique genes exhibited rates comparable to those of single-copy genes (Figure 1A). Thus, deleted unique genes generally do not appear to evolve under relaxed constraint or positive selection, instead showing levels of constraint similar to single-copy genes.

**Figure 1.**
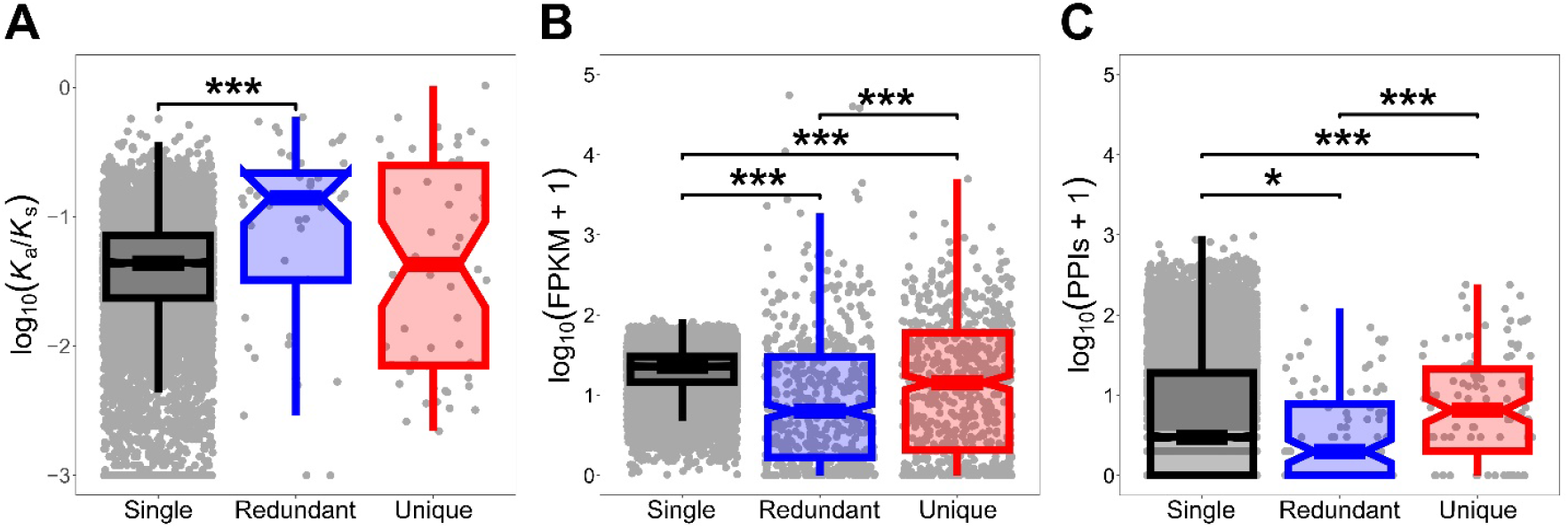
Sequence, expression, and protein-level metrics of deleted genes in *Drosophila*. Boxplots showing the distributions of (A) sequence evolutionary rates measured as log_10_(*K*_a_/*K*_s_), (B) gene expression levels given as log_10_(FPKM + 1), and (C) numbers of PPIs given as log_10_(PPIs + 1) for single-copy, deleted redundant, and deleted unique genes. \**p* < 0·05, \*\**p* < 0·01, \*\*\**p* < 0·001 (see *Methods*).

To gain further insight into this finding, we examined gene expression levels and numbers of protein-protein interactions (PPIs) of deleted genes, again using single-copy genes as a baseline.

Both deleted redundant and deleted unique genes had significantly lower expression than single-copy genes, with deleted unique genes showing significantly higher expression than deleted redundant genes (Figure 1B). Analyses of PPIs revealed contrasting patterns: deleted redundant genes participate in fewer interactions than single-copy genes, whereas deleted unique genes engage in more interactions than both single-copy and deleted redundant genes (Figure 1C). Taken together, these findings indicate that deleted unique children exhibit characteristics consistent with retained functionality.

Last, we leveraged recent advances in protein structure prediction (Jänes & Beltrao 2024) to compare the structural properties of single-copy, deleted redundant, and deleted unique genes. Predicted protein structures were obtained from AlphaFold (Jumper et al. 2021; Varadi et al. 2022, 2024) and aligned to compute TM-scores (see *Methods*), which range from 0 when structures show no similarity to 1 when they are identical (Zhang and Skolnick 2004). We did not detect statistically significant differences in TM-scores among gene categories (Figure 2A), potentially reflecting limited statistical power. We therefore also examined structural predicted local distance difference test (pLDDT) scores, which range from 0 to 100 and quantify confidence in predicted protein structures, with lower values typically associated with greater intrinsic disorder (Jumper et al. 2021). In this analysis, both deleted redundant and deleted unique genes generally exhibited higher pLDDT scores than single-copy genes (Figure 2B), indicating that deleted genes tend to be more structurally ordered than single-copy genes. Structural superimpositions further showed that, in both classes, the least similar deleted genes tended to be shorter and structurally simpler than the most similar deleted genes (Figure 2C-F).

**Figure 2.**
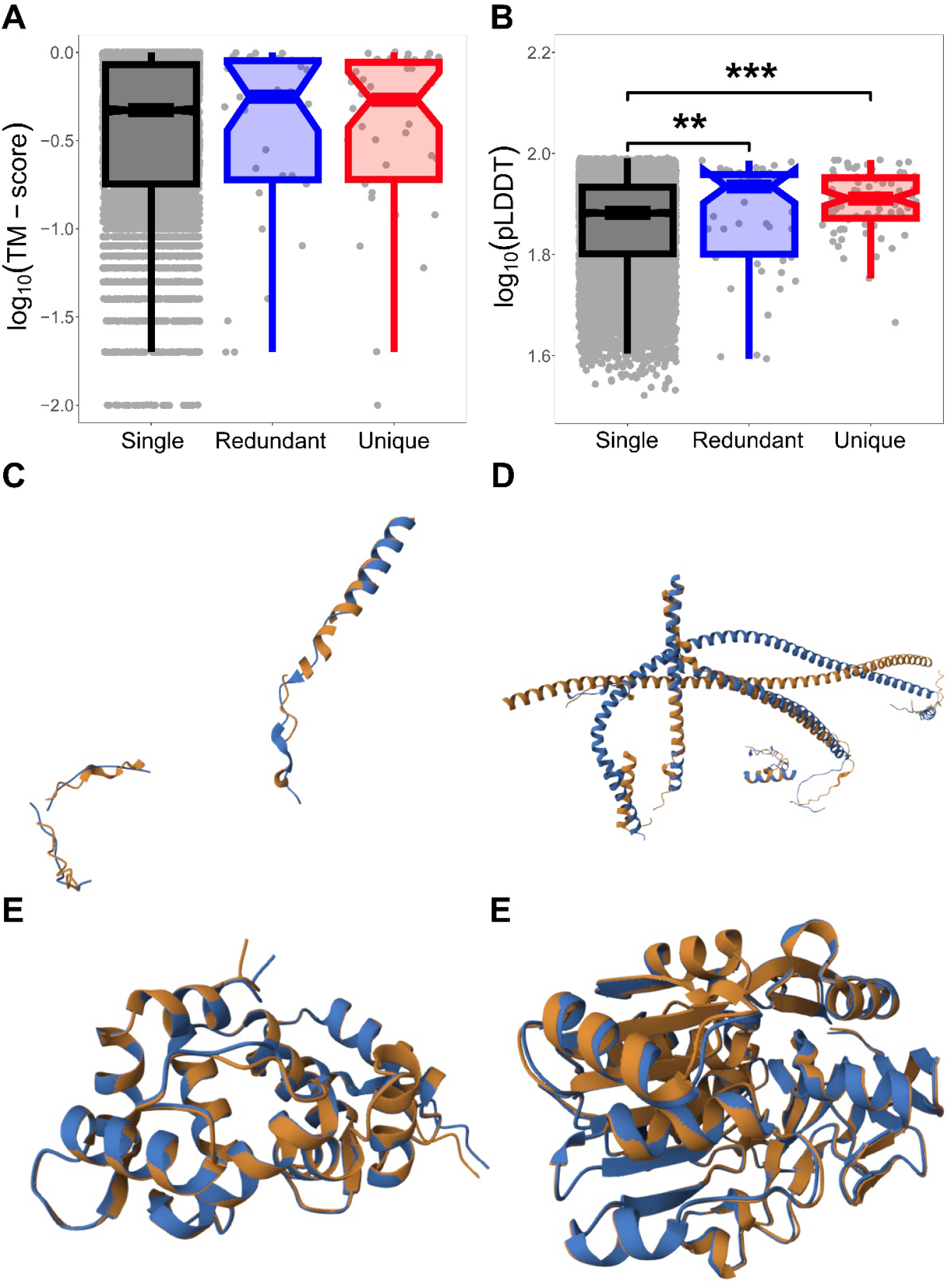
Protein structural properties of deleted genes in *Drosophila*. Boxplots showing the distributions of (A) protein structural similarity measured by log_10_(TM-scores) and (B) protein intrinsic disorder given as log_10_(pLDDT) scores for single-copy, deleted redundant, and deleted unique genes. (C-F) Structural superimpositions of deleted genes illustrating the lowest and highest structural similarity: (C) least similar deleted redundant genes, (D) least similar deleted unique genes, (E) most similar deleted redundant genes, and (F) most similar deleted unique genes. \**p* < 0·05, \*\**p* < 0·01, \*\*\**p* < 0·001 (see *Methods*).

## DISCUSSION

Gene deletion plays a key role in shaping genome architecture and evolution, yet the factors driving this process remain poorly understood. In this study, we systematically characterized 100 gene deletion events in *Drosophila* to uncover the evolutionary and functional patterns underlying gene loss. By integrating sequence, expression, interaction, and structural data, we identified distinct molecular signatures that distinguish functionally redundant and unique deletions. Together, these results provide insight into how gene deletion contributes to the refinement of genome function and the evolution of novel traits.

Our analyses revealed that child copies are preferentially deleted when duplicate genes have unique functions, whereas no such bias occurs for redundant duplicates. This finding suggests that the loss of unique child genes is not random but may reflect their transient evolutionary status. Two main hypotheses could explain this pattern. First, unique child genes may possess nonadaptive or weakly adaptive functions that are not maintained by selection, leading to relaxed constraint and eventual loss through drift. Alternatively, they may be actively purged by adaptive pressures that favor genome streamlining or the stabilization of regulatory networks.

The first hypothesis predicts that deleted unique genes should exhibit signatures of relaxed constraint, such as elevated *K*_a_/*K*_s_ ratios, low expression, limited protein-protein connectivity, and increased structural divergence (Kaessmann 2010; Assis & Bachtrog 2013; Kuzmin et al. 2022; Birchler & Yang 2022), consistent with the characteristics of young duplicates in *Drosophila* (Assis and Bachtrog 2013). However, our results do not support this model. Unique deleted genes show *K*_a_/*K*_s_ ratios comparable to those of single-copy genes and lower than redundant duplicates, indicating stronger selective constraint rather than relaxation. Likewise, their higher expression, greater network connectivity, and comparable structural divergence suggest that they remain functionally active prior to deletion. Additionally, these properties place deleted unique genes closer to single-copy genes than to redundant duplicates.

Because deletion primarily targets unique genes in *Drosophila* (Assis 2019; Campelo dos Santos et al. 2024), these findings challenge the expectation that weakly expressed, low-connectivity genes are the most dispensable (DeLuna et al. 2010; Birchler & Yang 2022). Instead, the observed properties may reflect a transient phase of integration into complex regulatory networks before removal, similar to patterns observed in other systems where young genes temporarily acquire extensive interactions before being lost (Li et al. 2005; Zhang et al. 2005; Jin et al. 2013). This dynamic suggests that gene deletion may contribute to the fine-tuning of molecular networks by adjusting connectivity and expression balance rather than merely eliminating redundancy. Collectively, these observations support the second model, in which adaptive purging removes transiently functional duplicates whose contributions are no longer advantageous, possibly reflecting selective pressures toward genome and regulatory streamlining.

Overall, our findings suggest that gene deletion in *Drosophila* is not a purely degenerative process but a dynamic force in genome evolution. The preferential loss of unique child genes, despite their functional activity, highlights the selective balance between innovation and constraint: while gene duplication promotes evolutionary novelty, deletion acts as a counterforce that removes transient or maladaptive innovations. This interplay likely maintains genome efficiency and regulatory stability while permitting bursts of evolutionary experimentation. Future comparative and functional studies will be critical to determine whether similar patterns of adaptive gene loss occur across taxa and to elucidate the mechanisms by which deletion shapes the long-term evolution of gene networks.

## METHODS

### Data acquisition and processing

The 100 deletion events analyzed in this study were obtained from the Dryad dataset associated with a previous study (Assis 2019; available at https://doi.org/10.5061/dryad.742564m). In that study, deletions were inferred to have occurred in either *D. melanogaster* or *D. pseudoobscura* after their divergence, based on phylogenetic comparisons across 12 fully sequenced and annotated *Drosophila* species (Drosophila 12 Genomes Consortium et al. 2007). The dataset also included 9,177 single-copy genes used as a comparative baseline, along with quantile-normalized expression levels in six tissues (carcass, male head, female head, testis, accessory gland, and ovary) for both single-copy and deleted genes. For all pairwise comparisons of sequence, expression, PPI, and structural features involving deleted genes, we analyzed the S and L copies, which represent orthologous genes between the two species. Summaries of all findings for deleted genes are presented in Table S1.

We supplemented the expression data from Assis (2019) with sequence, PPI, and protein structure information from additional sources. Coding sequences for all genes used in this study were downloaded from FlyBase (Öztürk-Çolak et al. 2024), and the longest transcript of each gene was retained. Sequences were aligned using MACSE v2 (Ranwez et al. 2018), and *K*_a_/*K*_s_ values were estimated with the codeml program in PAML (Yang 2007). PPI data were obtained from the STRING database (Szklarczyk et al. 2023). Analyses were restricted to physical links between proteins, and STRING alias files were used to identify the corresponding protein product for each gene. Predicted protein structures and associated pLDDT scores were obtained from AlphaFold v3 (Jumper et al. 2021; Varadi et al. 2024, 2022), and structural similarity was quantified using the template modeling (TM) score (Zhang & Skolnick 2004) via the Pairwise Structure Alignment tool (Bittrich et al. 2024) at the Protein Data Bank (Berman et al. 2000) with the jFATCAT (flexible) algorithm (Ye & Godzik 2003).

### Assignment of parent–child relationships

We began by reviewing the literature for each S and L gene to identify experimental evidence of their duplication history. Definitive parent***–***child relationships were found for only 10 of the 100 deletion events, all involving deletions in the *D. pseudoobscura* lineage such that both copies were present in *D. melanogaster*. This bias reflects the greater number of studies available for *D. melanogaster* relative to *D. pseudoobscura*.

Because the literature provided evidence for only a small subset of cases, we next examined additional features distinguishing S and L genes. First, we assessed sequence conservation across 12 *Drosophila* species (Drosophila 12 Genomes Consortium et al. 2007), under the expectation that parents should be more conserved than children. Using whole-genome alignments from the UCSC Genome Browser (Nassar et al. 2023) and the flyDIVaS database (Stanley & Kulathinal 2016), we designated the copy conserved across all 12 species as the parent, resolving 16 additional deletion events. We then examined gene structures for evidence of retrotransposition, in which an mRNA transcript is reverse-transcribed into cDNA and reinserted into the genome (Kaessmann et al. 2009). Retrotransposed duplicates are typically intronless, in contrast to their multi-exon ancestral genes (Langille & Clark 2007). Among the remaining 74 deletion events, we identified 22 in which one gene lacked introns while its paralog contained at least one, allowing us to designate the intronless copy as the child.

To further evaluate conservation, we queried OrthoDB v11 (Kuznetsov et al. 2023) for orthologs of the remaining S and L genes across 37 *Drosophila* species, designating the copy with more orthologs as the parent, consistent with greater evolutionary age. This approach resolved 17 additional cases, including 14 with orthologs present in all 37 species. We also reanalyzed the OrthoFinder (Emms & Kelly 2019) results from Assis (2019), which identified 21,944 orthogroups across 12 *Drosophila* species. For the unresolved events, we designated the copy belonging to the larger orthogroup as the parent, resolving 20 of the remaining 35 deletion events.

Finally, we applied the same conservation-based criteria to assess D genes in the remaining 15 unresolved events. This analysis yielded definitive parent–child assignments for five additional cases, while the remaining ten lacked sufficient evidence and were left unclassified.

### Statistical analyses

All statistical analyses were performed in R (R Core Team 2021). Exact two-sided binomial tests were conducted using the binom.test function in the *stats* package (R Core Team 2021) to compare observed and expected frequencies of different subclasses of deleted genes. For the comparison of resolved parent–child relationships between redundant and unique deletions, we set the number of successes *x* = 37 to represent the number of resolved redundant deletions, the number of trials *n* = 45 to represent the total number of redundant deletions, and the probability of success *p* = 53/55 to represent the proportion resolved among unique deletions. For the comparison of proportions of deleted parents and children, we set *x* = 23 to represent the number of deleted children, *n* = 37 to represent the total number of redundant deletions with resolved parent–child relationships, and *p* = 0·5 to represent the null expectation of no bias. For the comparison of proportions of deleted parents and children among unique deletions, we set *x* = 44 to represent the number of deleted children, *n* = 53 to represent the total number of unique deletions with resolved parent–child relationships, and *p* = 0·5 to represent the null expectation of no bias.

Differences between distributions shown in Figures 1 and 2 were assessed with two-sided Mann-Whitney *U* tests implemented with the wilcox.test function in the *stats* package (R Core Team 2021).

## ACKNOWLEDGEMENTS

This work was supported by National Institutes of Health grant R35GM142438 and National Science Foundation grant DBI-2130666 to R.A.

